# Comparative genomics of clinical *Stenotrophomonas maltophilia* isolates reveals regions of diversity which correlate with colonization and persistence in vivo

**DOI:** 10.1101/2023.07.14.549068

**Authors:** Melissa S. McDaniel, Nicholas A. Sumpter, Natalie R. Lindgren, Caitlin E. Billiot, W. Edward Swords

## Abstract

*Stenotrophomonas maltophilia* is a Gram-negative emerging opportunistic pathogen often found in respiratory diseases such as cystic fibrosis (CF). Patients with CF experience lifelong polymicrobial infections of the respiratory mucosa. Our prior work showed that *P. aeruginosa* promotes persistence of *S. maltophilia* mouse respiratory infections. As is typical for environmental opportunistic pathogens, *S. maltophilia* has a large genome and a high degree of genetic diversity. In this study, we evaluated the genomic content of *S. maltophilia,* combining short and long read sequencing to construct complete genomes of 10 clinical isolates which were then compared with the larger phylogeny of *S. maltophilia* genomic sequence data, and compared colonization/persistence in vivo, alone and in coinfection with *P. aeruginosa*. We found that while the overall genome size and GC content were fairly consistent, there was considerable variability in arrangement and gene content. Similarly, there was significant variability in *S. maltophilia* colonization and persistence in vivo in experimental mouse respiratory infection. Ultimately, this study gives us a greater understanding of the genomic diversity of *S. maltophilia* isolated from patients, and how this genomic diversity relates to interactions with other pulmonary pathogens, and to host disease progression. Identifying the molecular determinants of infection with *S. maltophilia* can facilitate development of novel antimicrobial strategies for a highly drug-resistant pathogen.

## INTRODUCTION

*Stenotrophomonas maltophilia* is a gram-negative bacterium that is widely distributed in environmental reservoirs such as water and soil, and is an emerging opportunistic pathogen in persons with cystic fibrosis (CF), particularly in the later stages of disease (1, 2). In the context of cystic fibrosis, defects in the cystic fibrosis transmembrane conductance regulator (CFTR) results in dehydrated and viscous mucus, a decrease in mucociliary clearance, and chronic inflammation. As a result, pwCF typically experience lifelong opportunistic airway infections that are polymicrobial in nature (3–9). Further, the delayed clearance and obstruction of airways results in a CF-specific phenomenon of micro-niche development in which regional adaptation in the colonizing microbial population over time leads to distinct populations within each bacterial species (10–14). Adaptation to the lung environment is very important for bacterial persistence, with specific factors required for initial colonization and others highly associated with chronic infection (15). In the case of *P. aeruginosa*, variants typically have changes in factors associated with motility, secreted toxins and proteases (16).

*S. maltophilia* is primarily environmentally acquired, and there is little evidence that strains are effectively transmitted between patients. While some human-infection associated factors have been identified, isolates must primarily rely on factors that also promote their survival in environmental situations to colonize the human lung (2, 17–21). Our previous work described a cooperative interaction between *S. maltophilia* and *P. aeruginosa* where damage to the lung epithelium by *P. aeruginosa* contributed to the increased persistence of *S. maltophilia* (22–24). These studies were, however, restricted to only a few strains, and given the amount of genetic diversity present in *S. maltophilia*, it is likely that genetic content plays a role in determining both the ability of an isolate to successfully infect a human host, and the amount of cooperativity that is seen with *P. aeruginosa*.

In this study, we sequenced the genomes of 10 clinical isolates of *S. maltophilia* from two different medical centers and evaluated the amount of genetic diversity present between these strains. The primary goal of this study was to look at the impact of genetic content on cooperativity that we see between *S. maltophilia* and *P. aeruginosa*, and on the ability of *S. maltophilia* to persist in the lung on its own. We found that although strains were similar in genomic size and percent GC content, there was significant genomic diversity between our strains, with large-scale rearrangements, insertions, and deletions when compared to the genome of *S. maltophilia* K279a. Strains were grouped into previously described phylogenetic clusters for *S. maltophilia.* Clustering did not correlate with medical center of isolation, or with survival of *S. maltophilia* in static biofilm coculture with *P. aeruginosa.* However, these clusters did correlate with both ability to persist in the murine lung alone, and the cooperative phenotype seen with *P. aeruginosa.* In this study we further characterized the genetic diversity seen within *S. maltophilia*, which enabled us to investigate the contribution of genomic content to persistence of *S. maltophilia* strains in the lung, both in the presence and absence of *P. aeruginosa*.

## RESULTS

To investigate the genetic diversity of *S. maltophilia* clinical isolates, we performed whole genome sequencing via a combination of short-read (Illumina) and long-read (Oxford Nanopore) sequencing on 10 clinical isolates from 2 medical centers. Hybrid assembly via Unicycler allowed us to construct the genomes with only a few contiguous segments each, ranging from 3 to 14 total contigs and an average L50 of 2.1. To complete the genomes, contigs were then reordered according to homology with *S. maltophilia* K279a using the Mauve Contig Mover. We found relatively little variation in both GC content and total genome length between strains, with GC content ranging from 66.2% to 67.4% and genome length ranging from ∼4.0kb to ∼4.8kb (Table 1). However, alignment of the constructed genomes revealed major rearrangements in comparison to *S. maltophilia* K279a. This included insertions and deletion, inversions, and large-scale movement of genetic islands (Figure 1).

**Figure 1.**
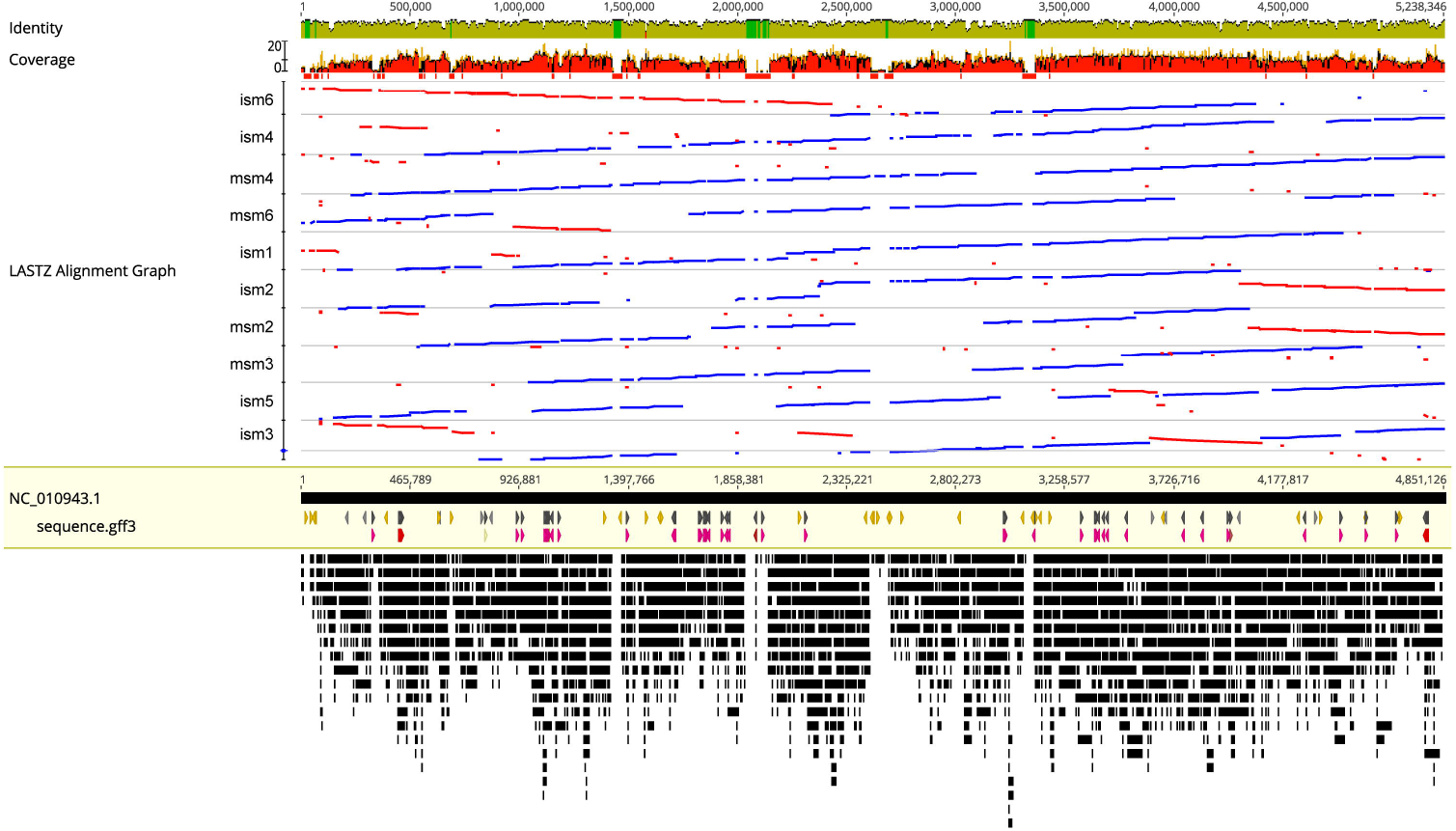
Alignment of clinical isolates to the genome of *S. maltophilia* K279a. Genomes of clinical isolates of *S. maltophilia* we sequenced using a combination of short read (Illumina) and long read (Oxford Nanopore) sequencing. Genomes were assembled via hybrid assembly using Unicycler, and contigs were rearranged according to the *S. maltophilia* K279a genome using Mauve Contig Mover. Whole genome alignments were performed using LastZ and visualized in Geneious. Blue lines represent regions of the genome in the same orientation as K279a, while red lines represent regions that are in the reverse orientation. Genomes are arranged in order of phylogeny.

**Table 1.** Summary statistics of genome assembly. *S. maltophilia* isolates were whole genome sequenced via a combination of short-read (Illumina) and long-read (Oxford Nanopore) sequencing. Genomes were assembled via hybrid assembly with Unicycler. Assembly quality was assessed using Quast, and gene annotation was performed with two commonly used annotation softwares, Prokka and RASTtk.

|  | Percent of Strains | Genes |
| --- | --- | --- |
| Core genes | 95 - 100 | 1209 |
| Soft core genes | 94 - 95 | 0 |
| Shell genes | 15 - 94 | 5037 |
| Cloud genes | 0 - 15 | 8313 |
| Total genes | 0 - 100 | 14559 |

**Table 1**
|  | Percent of Strains |
| --- | --- |
| Core genes | 95 - 100 |
| Soft core genes | 94 - 95 |
| Shell genes | 15 - 94 |
| Cloud genes | 0 - 15 |
| Total genes | 0 - 100 |

After assembling and annotating the genomes of our isolates, we then identified the core genome of *S. maltophilia* via Roary. We first performed this for only our sequenced strains as compared to *S. maltophilia* K279a. We found that among these strains, 1209 genes were found in at least 95% of the strains. In total, 14,559 genes were represented in the pangenome. We then performed this again using every published *S. maltophilia* genome, excluding those whose 16S internal transcribed spacer (ITS) sequence identified them as other *Stenotrophomonas* species, for a total of 372 strains. In this case we found a smaller core genome shared through the entire species, with 653 genes found in at least 95% of the strains annotated. As expected, the pangenome had a large number of total genes represented, with 87,043 identified (Table 2).

**Table 2.** Core genome assignment of *S. maltophilia.* Genomes of *S. maltophilia* from this study, and from the NCBI collection of published genomes were annotated via Prokka. The core genome of *S. maltophilia* was then assigned by identifying those genes shared by at least 95% of strains either in this study alone, or in the entire genome collection using Roary. The pangenome in each case was also calculated by adding the total number of unique genes represented by all strains.

|  | Medical center | GC% | Length | Contig # | CDS (prokka) | CDS (rastk) |
| --- | --- | --- | --- | --- | --- | --- |
| msm2 | UAB | 67.3 | 4466463 | 6 | 3966 | 4071 |
| msm3 | UAB | 66.7 | 4319629 | 5 | 3838 | 3948 |
| msm4 | UAB | 66.2 | 4756934 | 8 | 4375 | 4504 |
| msm6 | UAB | 66.7 | 4495881 | 6 | 3989 | 4115 |
| ism1 | IMC | 67.4 | 4449333 | 9 | 3996 | 4112 |
| ism2 | IMC | 67.4 | 4584707 | 10 | 4100 | 4229 |
| ism3 | IMC | 66.6 | 4611932 | 7 | 4188 | 4312 |
| ism4 | IMC | 66.7 | 4711422 | 14 | 4202 | 4317 |
| ism5 | IMC | 66.7 | 4591791 | 11 | 4128 | 4243 |
| ism6 | IMC | 67 | 3967727 | 3 | 3526 | 3622 |

Once a core genome was assigned, a core genome alignment for all published strains of *S. maltophilia*, along with the strains introduced in this study, were used to construct a phylogenetic tree (Figure 2). In previous work, an *in silico* multi locus sequencing type (MLST) scheme had been used to identify distinct phylogenetic clusters of *S. maltophilia*, many of which correlated with site of isolation (17). We identified most of these clusters in our tree, with 15 of the identified 23 represented. We found that our strains localized into a total of 6 of these clusters, with 2 of our strains found in branches that have not yet been defined. *S. maltophilia* strains ism3 and ism5 clustered with K279a into Sm6, a human respiratory associated cluster. Ism1, ism2, and msm2 clustered into Sm2, another human respiratory associated cluster. Msm4 and msm6 clustered into Sm3, a cluster predominantly made up of both human respiratory and human non-invasive strains. Ism4 was in cluster Sm12, which is primarily anthropogenic strains (associated with human environmental sources). The last two strains, ism6 and msm3 did not fit into any of the predefined clusters and were therefore assigned NC1 (new cluster 1) and NC2 (new cluster 2) respectively.

**Figure 2.**
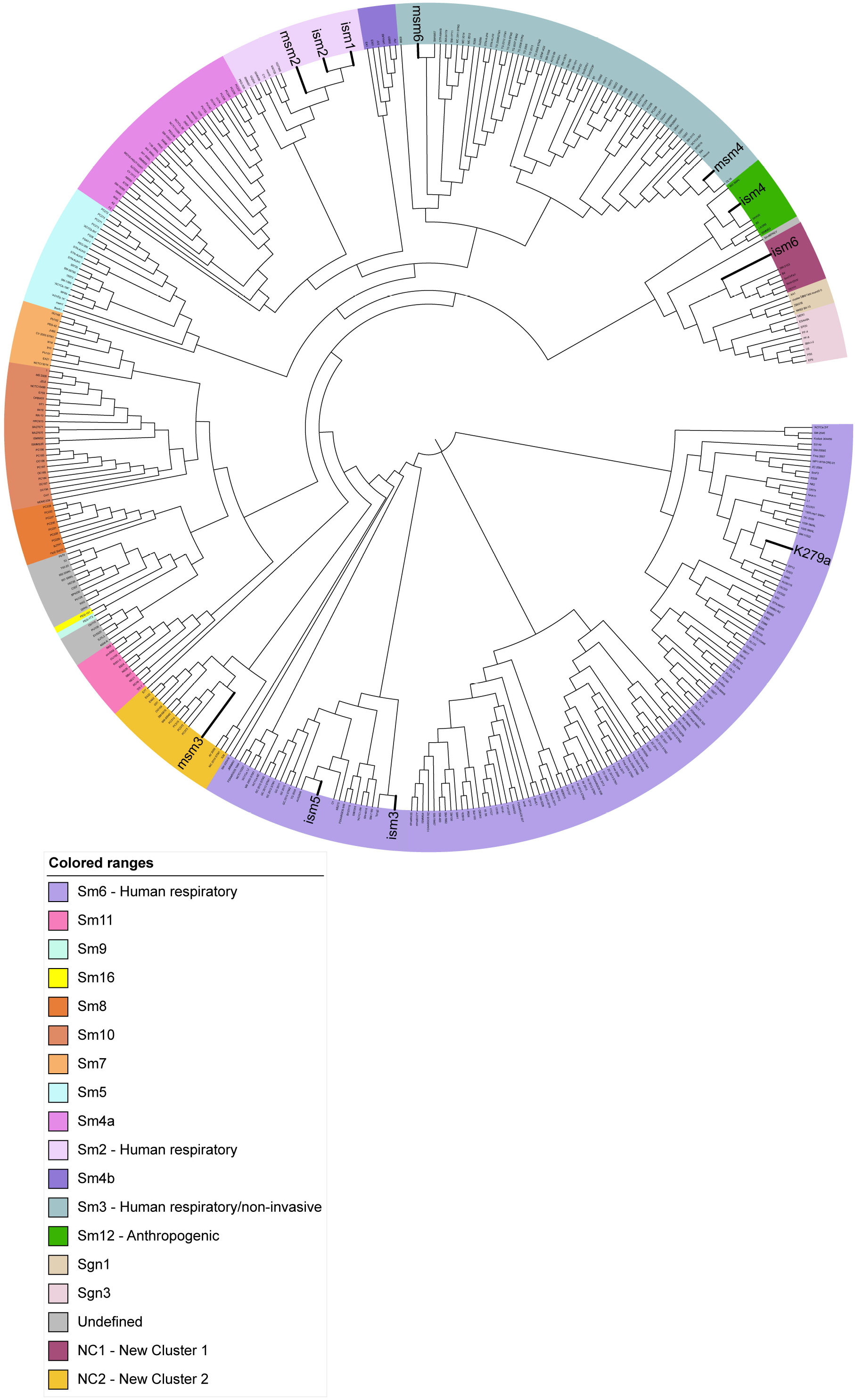
Phylogenetic tree of all published *S. maltophilia* isolates. A core genome alignment of all published isolates of *S. maltophilia*, along with those sequenced in this study, was used to construct a phylogenetic tree. Clusters were colored and labeled according to a previously published clustering scheme (10). Branch lengths were ignored for easier visualization.

To assess correlation of *S. maltophilia* phylogenetic relatedness with interaction in polymicrobial infections, we first measured the survival of clinical *S. maltophilia* isolates after coculture with *P. aeruginosa* in vitro. Although we have previously found that these organisms act cooperatively during pulmonary infections, there have been reports of antagonism between some strains during *in vitro* coculture (22, 25). We grew each clinical isolate of *S. maltophilia* in a polymicrobial biofilm with *P. aeruginosa* mPA08-31 at 30°C for 12 hours to assess its survival. We found that *S. maltophilia* strains showed a variable amount of inhibition in the presence of *P. aeruginosa,* but that this did not correlate with phylogenetic position (Figure 3A). *P. aeruginosa* counts were not affected by presence of any of the *S. maltophilia* isolates (Figure 3B).

**Figure 3.**
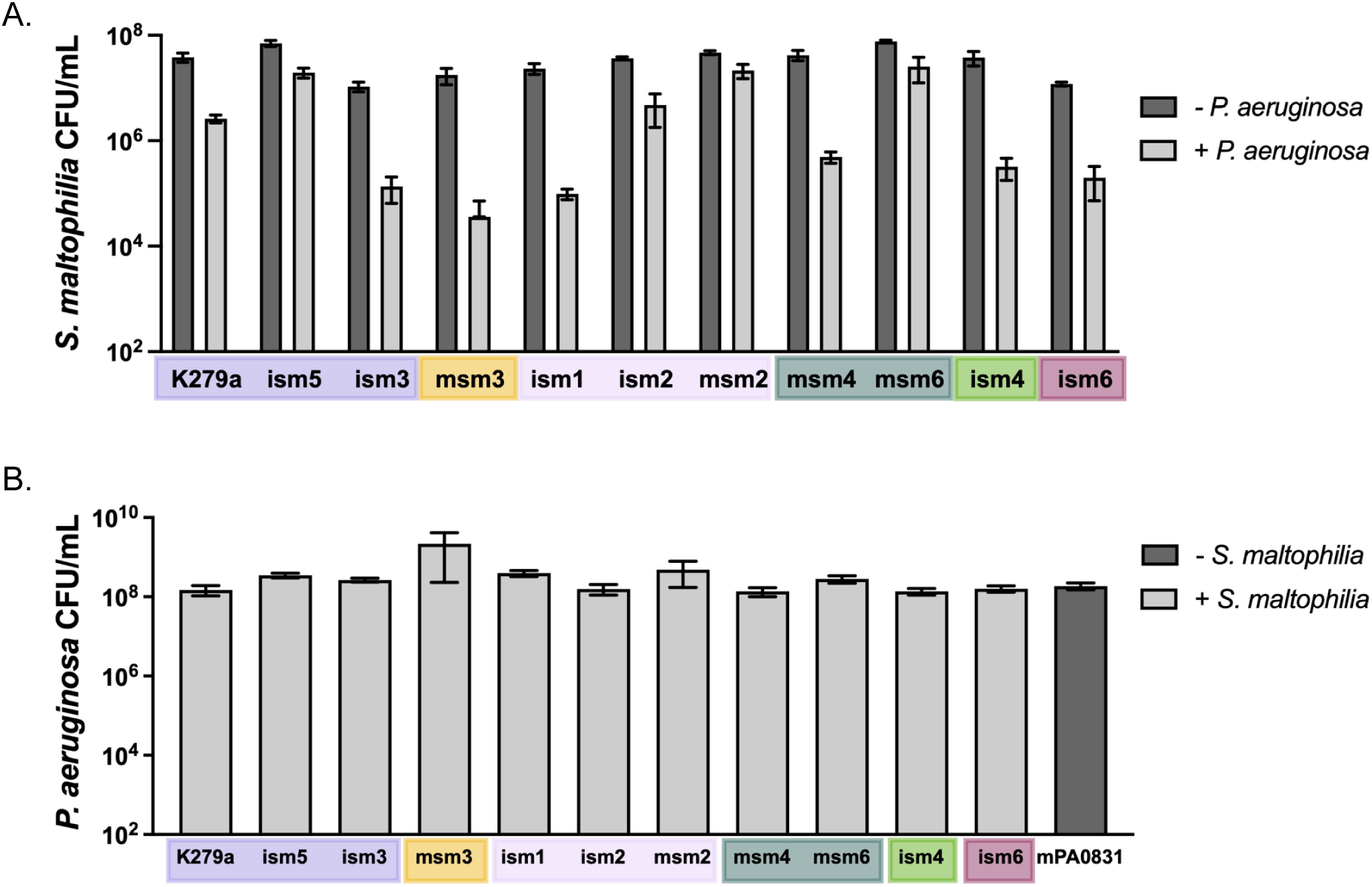
Survival of *S. maltophilia* in a static biofilm co-culture with *P. aeruginosa.* Static single- and dual-species biofilms of A) *S. maltophilia* K279a, ism1, ism2, ism3, ism4, ism5, ism6, msm2, msm3, msm4, msm6 and B) *P. aeruginosa* mPA0831 were seeded at ∼10 CFU/mL of each organism in LB and grown at 30°C for 12 hours. Mean ± SEM, n = 3.

With these data in hand, we next performed mouse respiratory infections with each strain individually, and in polymicrobial infection with *P. aeruginosa.* Unlike in the *in vitro* biofilms, we found that phylogenetic clusters did correlate with infection outcomes, both in single-species and in polymicrobial infections. While most strains had significantly higher *S. maltophilia* burden during polymicrobial infection, one phylogenetic cluster (Sm2) had no evidence of cooperativity, largely due to an increased colonization/persistence of strains of *S. maltophilia* in this cluster during single-species infection. The 2 strains that share a cluster with K279a, ism3 and ism5, both has a significantly higher burden of *S. maltophilia* during polymicrobial infection (*P* = 0.019 and *P* = 0.013 respectively) and had a similar burden to that of *S. maltophilia* K279a during single-species infection. Msm3, which clustered on its own, persisted to a lesser degree than *S. maltophilia* K279a, and trended toward a cooperative phenotype, though it was not statistically significant. Strain ism6, which also formed a new cluster, was the only isolate to persist to the same degree as *S. maltophilia* K279a but show no evidence of cooperativity during polymicrobial infection with *P. aeruginosa* (Figure 4A).

**Figure 4.**
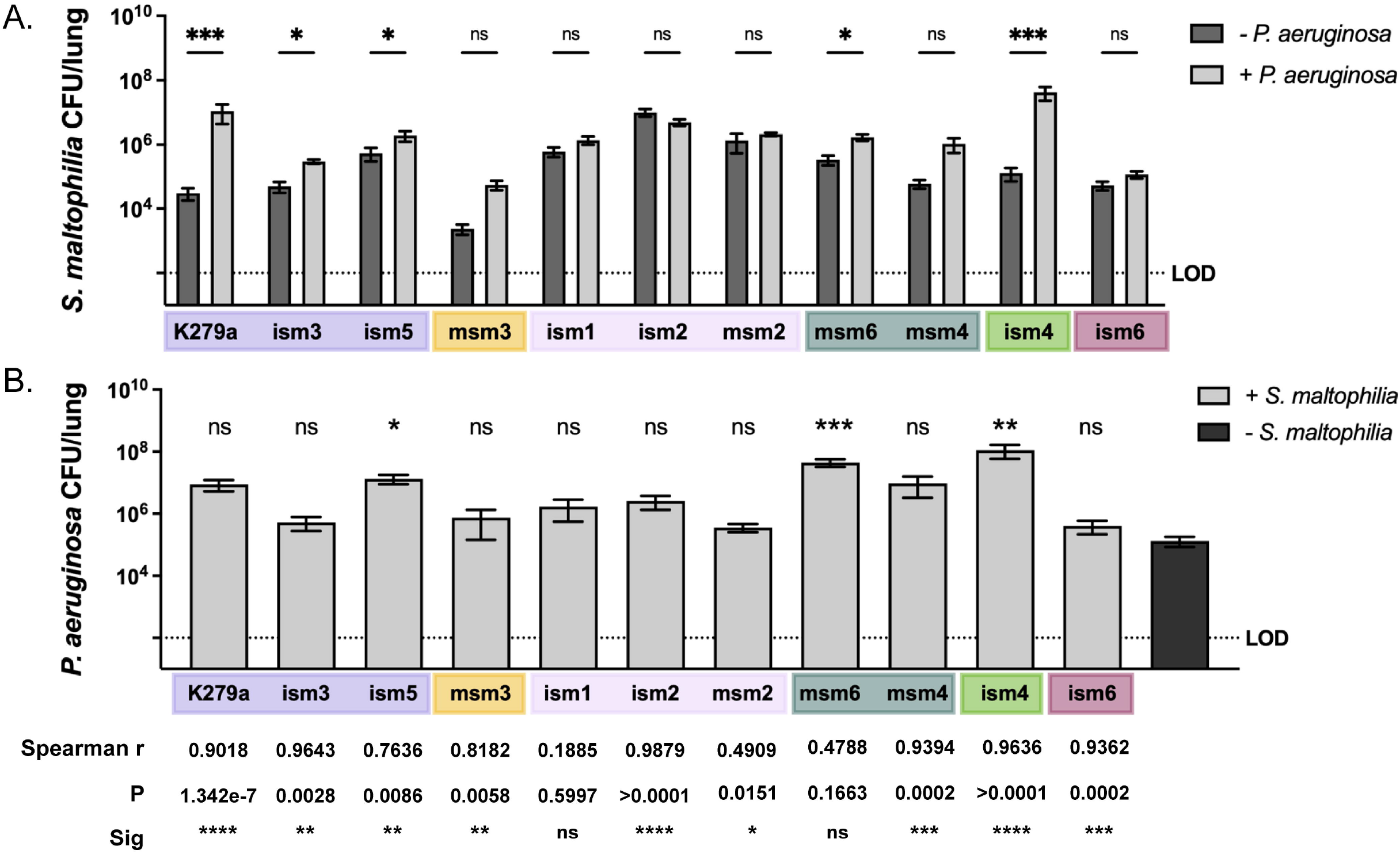
Persistence of *S. maltophilia* strains in the murine lung during polymicrobial infection with *P. aeruginosa.* BALB/cJ mice were intratracheally infected with ∼10^7^ CFU of *S. maltophilia* K279a, ism1, ism2, ism3, ism4, ism5, ism6, msm2, msm3, or msm4 and *P. aeruginosa* mPA0831 alone and in combination. Groups were euthanized at 24 hours post-infection. Bacterial load in lung homogenate was enumerated via viable colony counting on differential medium. Mean ± SEM, n = 7 to 9. Kruskal-Wallis test with uncorrected Dunn’s *post hoc* comparisons, *, *P* < 0.05; **, *P* < 0.01; ***, *P* < 0.001. Significant outliers were identified via ROUT method and removed. Colors correspond to phylogenetic clusters defined in Figure 1.

Consistent with previous studies, we found that polymicrobial infection with *S. maltophilia* K279a provided no benefit to the persistence of *P. aeruginosa.* However, this was not the case for all the clinical isolates tested. Polymicrobial infection with several *S. maltophilia* isolates, particularly those with the highest degree of cooperativity, produced an increase in the burden of *P. aeruginosa* in the lung (Figure 4B). Despite heterogeneity in the amount that these organisms were able to promote the bacterial burden of one another, nearly all the strains showed a correlation in the bacterial burden of *S. maltophilia* and *P. aeruginosa* in the lung, with strains ism3, ism2, and msm2 being the exceptions (Figure S1).

Our previous work investigating the mechanisms behind dual-species cooperativity between *S. maltophilia* and *P. aeruginosa* implicated *chpA*, the histidine kinase portion of a two-component regulatory system known to govern the type IV pilus in *P. aeruginosa*, as a determinant of the relationship between *S. maltophilia* and *P. aeruginosa* (26, 27). We therefore decided to explore the variability in this system seen in our clinical isolates. We identified the *chpA* locus via RASTtk annotation and homology to the locus in *S. maltophilia* K279a. We found that while the complete *chpA* locus was present in all of our strains, that there were many SNPs, insertions, and deletions present as compared to *S. maltophilia* K279a (Fig. 5A). Adherence assays on polarized cystic fibrosis bronchial epithelial cells (CFBEs) demonstrated binding patterns that differed based on phylogeny. *S. maltophilia* strains from the Sm2 cluster (ism1 and msm2) bound poorly to polarized epithelium, which did not change with prior infection with *P. aeruginosa. S. maltophilia* strains from Sm3 and Sm12 clusters (msm4 and ism4 respectively) bound more efficiently to the epithelial layer, which was slightly increased with prior infection with *P. aeruginosa,* although not to a statistically significant degree (Fig. 5B). To investigate the overall relationship of our clinical strains in relation to the *chpA* locus, we constructed a phylogeny based on the alignment of the *chpA* locus and found that phylogeny mirrors the relationships determined by the core genome (Fig. 5C).

**Figure 5.**
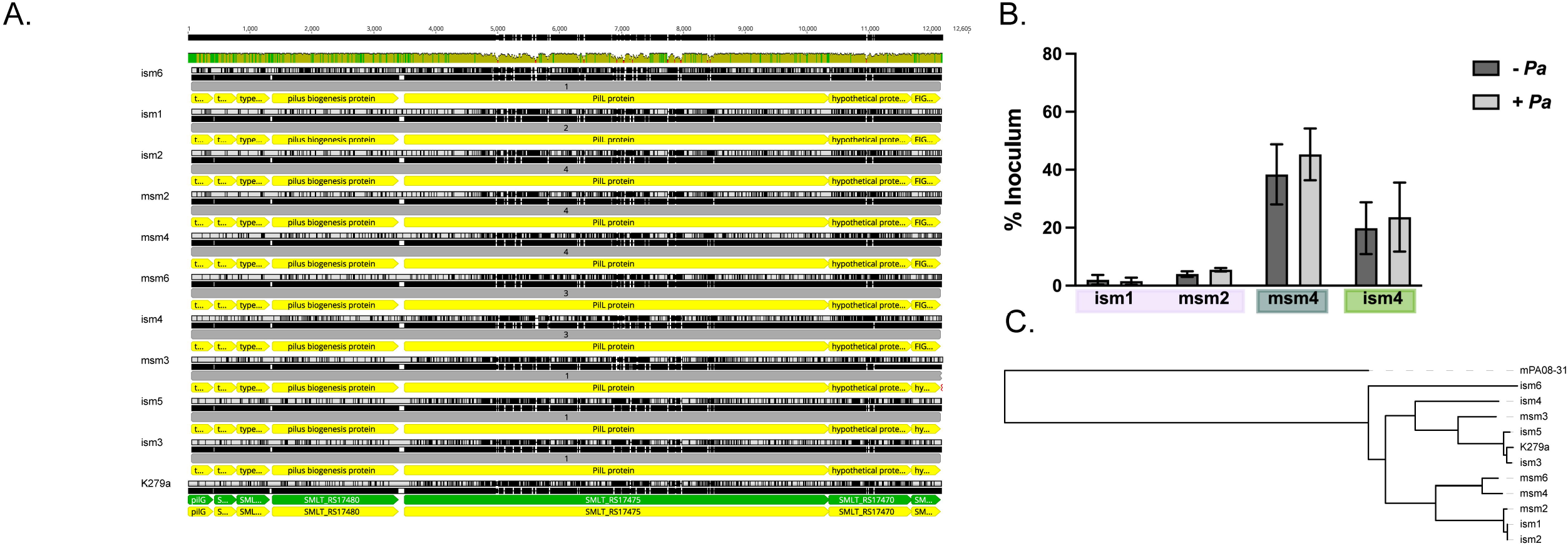
Analysis of variation in the *chpA* locus of *S. maltophilia*. The *chpA* loci of *S. maltophilia* clinical isolates were identified via annotation with RASTtk and homology to *S. maltophilia* K279a. A) Sequences were aligned via CLUSTALW on Geneious. B) Polarized CFBEs were pre-treated with either MEM or ∼10^6^ CFU of *P. aeruginosa* mPA08-31 for 4 hours before the addition of ∼10^6^ *S. maltophilia* ism1, ism2, msm4, or ism4. B) Results expressed as percent of initial *S. maltophilia* inoculum (Mean ± SEM, n = 9 wells). C) Phylogenetic relationship between the *chpA* locus of clinical strains of *S. maltophilia*, with *P. aeruginosa* mPA08-31 represented as an outgroup.

## DISCUSSION

In this study, we investigated the genetic diversity of clinical *S. maltophilia* strains and their ability to persist in the murine lung, alone and in combination with the important pulmonary pathogen *P. aeruginosa.* To do this, we used a combination of long and short read sequencing to assemble genomes to near completion, before comparing gene content and assessing their position within the larger phylogeny of *S. maltophilia*. We found that phylogenetic position of *S. maltophilia* strains correlates with ability to persist in the murine lung, and with the amount of cooperativity that is seen during polymicrobial infection with *P. aeruginosa*.

This study highlights the importance of using long read sequencing in combination with short reads for improved contiguity, and thus improved ability to detect structural rearrangements and insertions/deletions. Many of the insertions/deletions seen had clear evidence of either phage genes or transposon genes, explaining the large-scale rearrangements seen between strains. This also indicates that *S. maltophilia* might easily acquire genes from the environment, letting them quickly adapt, and contributing to the transition from environmental to medical contexts.

Although we were able to achieve a relatively good read depth, this was still not sufficient to completely circularize the genome. This made it difficult to come to be certain about every rearrangement that we observed. Increasing the read depth of long-read sequencing might resolve this problem. However, most of the junctions between contiguous sections occurred at repetitive regions, or those with high GC content, which might not be surmountable with deeper sequencing alone. This highlights the difficulty of fully sequencing large and GC-rich genomes, like those of *S. maltophilia*.

Interestingly, the phylogenetic cluster that showed the least cooperativity with *P. aeruginosa* (Sm2) was able to persist in the murine lung more successfully on its own when compared to other strains. It also clustered with isolates commonly found in pulmonary infections. This was not true of the other pulmonary-associated cluster (Sm6), where all tested strains showed significant cooperativity with *P. aeruginosa.* It is possible that we are seeing two different populations, one isolated from the lung early in infection, and one that has already adapted to the lung over time. The inverse relationship between adherence to epithelial cells and persistence in the animals could be indicative of adaptation to the lung, with factors like motility being some of the first that are lost in chronic infection.

We were able to correlate our strains to phylogenetic clusters already associated with specific sites of origin, giving us insight into the genetic factors required for persistence in different disease states. We were, unfortunately, missing specific metadata associated with most of our samples, including site of isolation, disease severity metrics, and information on co-isolation with other organisms. Performing a similar study on a larger number of samples, correlated with patient outcomes would allow us to dive deeper into the molecular determinants important in successful *S. maltophilia* infections, and those that govern co-infection with other pulmonary pathogens.

Taken together, the results of this study indicate that there is significant diversity within *S. maltophilia* isolates that infect humans, and that this genetic content correlates with ability to successfully colonize the mammalian lung. It also indicates that while lung-adapted *S. maltophilia* might not need help from external sources to persist, that non-lung adapted *S. maltophilia* strains may cooperate with other pulmonary pathogens like *P. aeruginosa* to cause disease. These results prompt further study into the specific genomic content of *S. maltophilia* that contributes to worst patient outcomes, and disease progression.

## METHODS

### Strains and growth conditions

*S. maltophilia* K279a is a widely used model strain, and its genome has been fully annotated and sequenced (28); this strain was provided by M. Herman (Kansas State University). *S. maltophilia* msm2, msm3, msm4, and msm6 are clinical isolates from patients at the UAB Medical Center in Birmingham, Alabama and were provided by W. Benjamin (University of Alabama at Birmingham). *S. maltophilia* ism1, ism2, ism3, ism4, ism5, and ism6 are clinical isolates from patients at UI Health Care in Iowa City, Iowa and were provided by T. Starner (University of Wisconsin-Madison). *P. aeruginosa* PAO1 was provided by D. Wozniak (Ohio State University), *P. aeruginosa* mPA08-31 was obtained from S. Birket (University of Alabama at Birmingham). All strains were routinely cultured on Luria Bertani (LB) agar (Difco) or LB broth. *S. maltophilia* strains were streaked for colony isolation before inoculating into LB broth and shaking overnight at 30°C, 200rpm. *P. aeruginosa* strains were streaked for colony isolation before inoculating into LB broth and shaking overnight at 37°C, 200 rpm.

### DNA extraction and sequencing

Overnight broth cultures of each strain were prepared in LB broth from a single colony to ensure a clonal population. Cultures were grown shaking overnight at 30°C, 200rpm. Genomic DNA was extracted using the DNeasy Blood and Tissue kit (Qiagen) with modifications for Gram-negative bacteria. Preparations were quantified via nanodrop, and quality of the preparation (size of fragments and specificity for DNA) was checked via gel electrophoresis before sequencing.

### Genomic assembly and annotation

For short read Illumina sequencing, samples were sent to the microbial genome sequencing center (MIGS) (Pittsburgh, PA). DNA sequencing was performed using an Illumina NextSeq2000 to generate paired end 151bp reads, with ∼3,000,000 reads per sample. For long read sequencing, Oxford Nanopore sequencing was also performed through MIGS, with ∼250,000 long reads per sample, with an average length of ∼10kb each. Reads were trimmed with bcl2fastq (v. 2.20.0.445, short reads) and Porechop (v. 0.2.3_seqan2.1.1, long reads) before genome assembly and annotation. Genomes were assembled *de novo* using hybrid short and long read assembly with Unicycler (v. 0.4.8). Quality of the assembly was evaluated via QUAST (v. 5.0.2) with *S. maltophilia* K279a as the reference genome. To finish assembly, multiple contigs were combined into a continuous genome using Mauve Contig Mover in Geneious (Auckland, New Zealand, Mauve plugin v. 1.1.1) with *S. maltophilia* K279a as a scaffold. Arranged contigs were then concatenated before moving to the annotation step. All genomes were annotated using Prokka (v. 1.14.5) and RASTtk (v. 1.3.0).

### Core genome determination and phylogenetic analysis

To evaluate the core genome of *S. maltophilia,* assembled sequences from all published *S. maltophilia* genomes were downloaded from NCBI and were annotated for gene content via Prokka. Genome content was compared using Roary, and the core genome was designated as genes present in at least 95% of the *S. maltophilia* strains. Phylogenetic relationships between strains were determined using an alignment of the core genome between strains. A tree was generated via FastTree (v. 2.1.12) and visualized in interactive tree of life (iTOL).

### Static biofilm assay

*In vitro* biofilm assays were performed according to an established micro-titer assay protocol (29) with some modifications. Biofilms were prepared from overnight broth cultures of *S. maltophilia* and *P. aeruginosa* diluted to an OD_600_ of 0.15 (∼10^8^ CFU/mL) in LB broth. Bacterial suspensions were used to inoculate a 96-well microtiter dish (200 μL/well) and biofilms were grown at or 37° C for 12 hours. Mature biofilms were washed twice with sterile PBS before plating on M9 minimal medium to select for *P. aeruginosa* and LB with gentamicin (50 µg/mL) to select for *S. maltophilia*.

### Mouse respiratory infections

BALB/cJ mice (8-10 weeks old) were obtained from Jackson laboratories (Bar Harbor, ME). Mice were anesthetized with isoflurane and intratracheally infected with either *S. maltophilia, P. aeruginosa,* or both (∼10^7^ CFU each in 100 µL PBS). The left lung of each mouse was harvested and homogenized in 500 µL of sterile PBS for viable plate counting. Homogenate from single-species infections were serially diluted in PBS and plated on LB to obtain viable CFU counts. Homogenate from polymicrobial infections were plated on M9 minimal medium (21) for *P. aeruginosa* and LB agar containing gentamicin (50 µg/mL) to enumerate *S. maltophilia.* All samples from polymicrobial infections were also plated on LB for total bacterial counts. All mouse infection protocols were approved by the UAB Institutional Animal Care and Use Committees.

### Ethical approval and compliance

All animal experiments were performed according to with ARRIVE international guidelines for animal research experimental protocols reviewed and fully approved by the UAB Animal Care and Use Committee and include detailed explanation of experimental design, randomization of experimental and control groups to ensure minimization of bias, and estimation of animal group sample size based on statistical power calculations to minimize numbers of animals necessary for valid statistical analyses. Anesthesia and euthanasia of mice were performed according to standards established by the American Veterinary Medical Association and under supervision of the UAB Animal Research Program.

### Cell culture

Cystic fibrosis bronchial epithelial cells (CFBE41o-) cells are an immortalized human bronchial epithelial cell line homologous for the F508del mutation in CFTR (30). Cells were routinely cultured in minimal essential medium (MEM, Corning) with 10% fetal bovine serum, and were polarized by seeding at a density of 5 x 10^6^ on the apical surface of transwells (0.4um, Corning) and growing at 37°C for 7 days, before removing the apical media and growing for an additional 7 days at air-liquid interface. Polarization of the epithelial membranes was confirmed via transepithelial electrical resistance measurements performed via EVOM^2^ Volt/Ohm Meter (World Precision Instruments).

### Adherence assays

To measure the adherence of *S. maltophilia* to CFBE41o-s after prior infection with *P. aeruginosa,* cells were inoculated with ∼10^6^ CFUs (MOI = 20) of *P. aeruginosa* mPA08-31 in MEM (FBS free). Bacteria were incubated on the cells for 4 hours before being removed from the apical chamber. Cells were then inoculated with ∼10^6^ CFUs (MOI = 20) of *S. maltophilia* (various strains) and incubated for an hour. Cells were washed twice with sterile 1X PBS before being scraped from the transwell membrane, diluted, and plated on differential medium to enumerate the bacterial burden.

### Statistical analyses

Unless otherwise noted, graphs represent sample means ± SEM. For non-parametric analyses, differences between groups were analyzed by Kruskal-Wallis test with the uncorrected Dunn’s test for multiple comparisons. For normally distributed data sets (as determined by Shapiro-Wilk normality test) a one-way ANOVA was used with Tukey’s multiple comparisons test. Outliers were detected via the ROUT method (Q=1%) and excluded from the analysis. All statistical tests were performed using Graphpad Prism 9 (San Diego, CA).

## ACKNOWLEDGMENTS

This work was supported by grants from the Cystic Fibrosis Foundation (CFFSWORDS1810) awarded to W.E.S. as well as NIH P30 center grant DK072482 and the Cystic Fibrosis Foundation Basic Research Center awarded to Dr. Steve Rowe. M.S.M. was supported by an NHBLI T32 UAB Predoctoral Training Program in Lung Diseases (T32HL134640) and as predoctoral fellow on the UAB Center for Cystic Fibrosis Research (P30 DK072482 and Cystic Fibrosis Foundation Research Development Program ROWE19RO).

The authors thank Dr. Bill Benjamin (University of Alabama at Birmingham), Dr. Susan Birket (The University of Alabama at Birmingham), and Dr. Timothy Starner (University of Wisconsin-Madison) for providing bacterial strains for this study.

